# Vascularized Organoids Emerge from a Heart–Foregut-like Intermediate Recapitulating Early Human Development

**DOI:** 10.64898/2026.10.21.758395

**Authors:** Wuming Wang, Qiqi He, Zhiqiang Xiong, Hong Zhang, Qi Meng, Xianwei Su, Ya Guo, Yong Fan, Luke Yunfei Yang, Xingguo Liu, Xiuwu Bian, Gang Lu, Wai-Yee Chan

**Author notes:** Address correspondence to (G.L.); (W.-Y. C.).

## Abstract

During early cardiogenesis, the heart regions lie in close contact with foregut endoderm, which supplies essential nutrients and signaling cues to orchestrate primitive heart tube assembly. Here, we show that self-organized heart/foregut like-structures emerge from human pluripotent stem cells (hPSCs). The initial heart/foregut–like structures, termed the “Seed”, are encased by surrounding supportive cells termed “Soil”, which provide a matrix microenvironment; the “Seed” progressively develops into chambered human cardiac organoids (hCOs) containing cardiomyocytes (CMs), epicardial cells, cardiac fibroblasts, and endothelial cells. Optimization of the induction system further enhanced hCO maturation and vascularization. Therefore, this “Soil–Seed” system provides a model to dissect human early heart/foregut self-organization.

## Introduction

During early embryonic development, the heart regions lie in close contact with foregut endoderm, and the foregut and the heart tube undergo coordinated elongation ^1^. The foregut endoderm and myocardium join forces to drive early heart tube assembly ^2^. Indeed, growth factors secreted by the nearby foregut endoderm promote the specification of cardiopulmonary progenitor ^3^. Therefore, studies of human early heart development should not only consider cardiac cell lineages, but also the foregut endoderm.

Human heart tube initiation remains poorly understood due to the lack of faithful experimental models. Although animal models are widely used, interspecies structural and genetic differences pose major barriers. Self-organizing three-dimensional (3D) cardiac organoids have advanced modeling of human cardiogenesis ^4,5^, and recent multi-chamber cardioid systems model first and second heart fields and early heart tube stages ^6^, yet these human cardiac organoids (hCOs) lack any foregut component or intermediate, and thus cannot comprehensively mimic human early cardiac development.

Drakhlis et al ^7^ reported a cardiac-foregut co-development model under the extracellular matrix or Matrigel conditions, tracing the early stages of human heart development. Co-emergent cardiac–foregut/gut tissues can progress toward heart tube formation ^8^. Distinct from previously reported cardiac-foregut models ^7^, we demonstrated that heart/foregut-like organoids can faithfully recapitulate human early heart tube formation.

## Results

### A self-organized heart/foregut-like intermediate serves as the “seed” for hCO induction

Human heart/foregut organoids can be generated from a 3D cultured system within extracellular matrix (ECM) microenvironments by modulating WNT signalling^7^. Here, we developed hCOs derived from heart/foregut-like intermediate structures (Fig. 1a). To monitor hCO formation, we stained the day-7 intermediate structures with anti-cTNT and anti-FOXA2 antibodies and observed numerous transient heart–foregut-like structures composed of cTNT-positive CMs and FOXA2-positive foregut endodermal cells appearing at day 7 (Fig. 1b, c). We hypothesize that this structure emerges briefly following WNT activation, reflecting a short-lived heart/foregut endoderm during *in vivo* early embryonic development ^7^. We propose that these conditions promote the formation of a heterogeneous multicellular population and a 3D microenvironment squeezed out by the densely packed cells at day 7 (Fig. 1a □ c), termed the “Soil”, which provides a supportive substrate for heart/foregut formation. We term these heart/foregut structures as “Seeds”, which have the potential to develop into miniature hCOs (Fig. 1a □ c), and this culture system was referred to as the “Soil–Seed” system.

**Figure 1.**
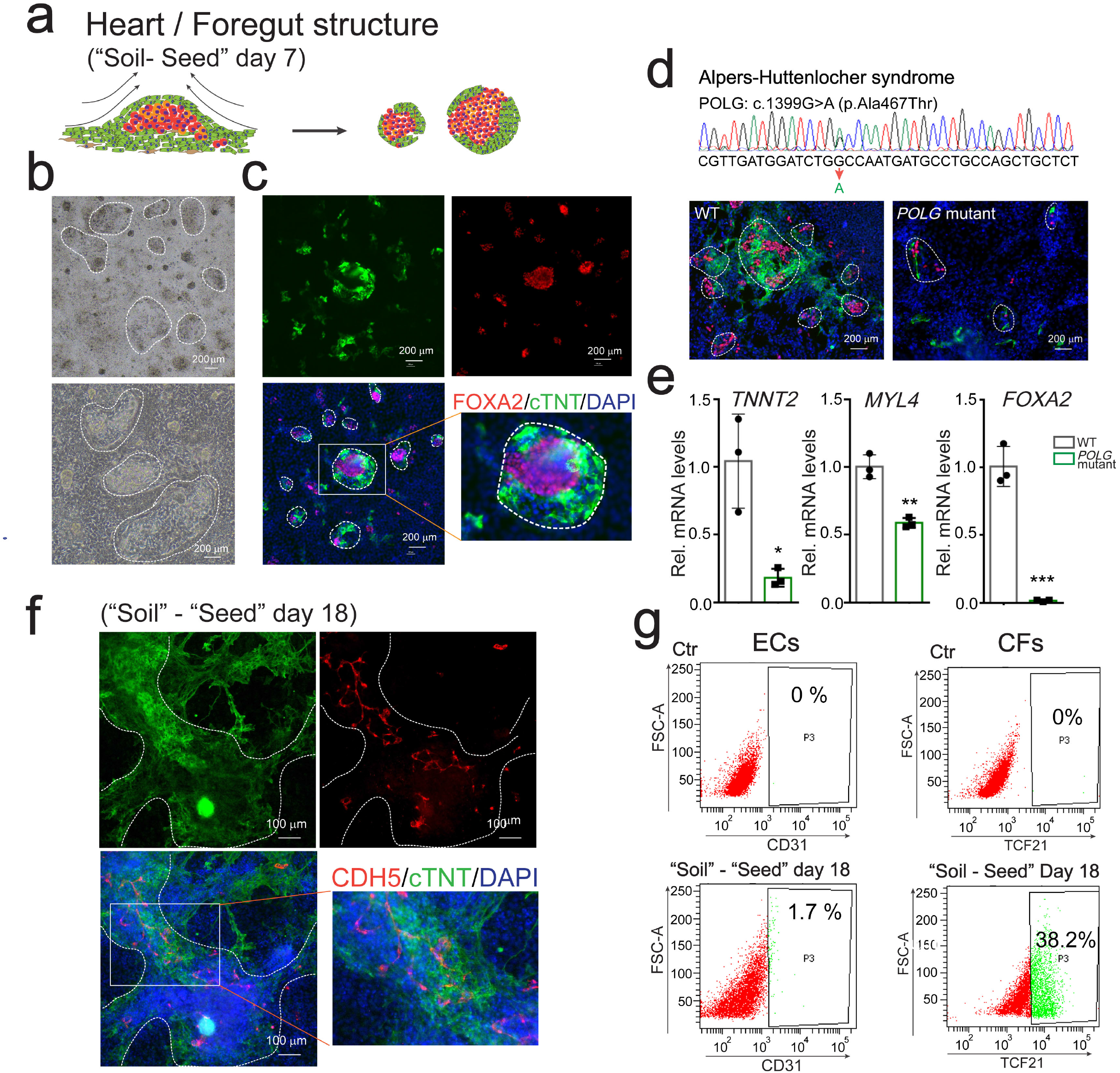
The “soil–seed” concept represents a foregut intermediate stage during hCO formation. **a**.Graphic showing the progress of “Soil – Seed” formation. **b**.Phase-contrast microscopy showing the grown out “Soil – Seed” at day 7. **c**.mmunofluorescence (IF) staining showing the heart/foregut assemble structure, also termed as “Seed”, including cTNT-positive CMs and FOXA2-positive foregut endoderm cells. **d**.iPSCs derived from an Alpers–Huttenlocher syndrome (AHS) patient carrying a POLG mutation (c.1399G>A) exhibit impaired heart and foregut formation at day 7. **e**.qPCR validation confirmed reduced expression of CM markers (*TNNT2, MYL4*) and the foregut marker (*FOXA2*) in AHS patient-derived cells harboring *POLG* mutations at day 7. **f**.IF visualization of aligned ECs and CMs within the “Soil” at day 18. **g**.Flow cytometry quantification showing approximately 1.7% CD31-positive ECs and 38.2% cardiac fibroblasts (CFs) in the “Soil” at day 18.

*POLG* mutations are the most common cause of inherited mitochondrial disorders, including Alpers–Huttenlocher syndrome (AHS), a multisystem condition affecting the brain, heart, liver, kidney, gastrointestinal tract, and respiratory system^9^. Given that the foregut is an embryonic region that gives rise to multiple organs—including the oesophagus, stomach, proximal duodenum, liver, pancreas, and parts of the respiratory tract—we hypothesized that the heart/foregut organoid system represents an ideal platform for studying AHS. Using AHS patient-derived iPSCs bearing a *POLG* mutation (c.1399G>A), we initiated heart/foregut formation and observed significant impairment in both heart/foregut development and expression of key genes, including cardiac genes (*TNNT2, MYL4*) and the foregut marker *FOXA2* (Fig. 1d, e).

By day 18, analysis of the “Soil” revealed vascular ECs growing in close association with CMs, as indicated by CDH5 expression (Fig. 1f). We propose that ECs and CMs both originate from mesodermal progenitors during early embryogenesis and engage in reciprocal crosstalk that drives their coordinated spatial organization. Flow cytometry confirmed the presence of 1.7% CD31-positive ECs and 38.2% TCF21-positive CFs (Fig. 1g). Together, these observations indicate that hCOs undergo self-organization from the “Soil,” initiating with a foregut-like intermediate that recapitulates key stages of human cardiac development and provides a relevant model for studying genetic variants associated with congenital embryonic defects.

### Single-cell transcriptomic profiling reveals cellular heterogeneity within hCOs and supporting “Soil”

To define cellular heterogeneity within hCOs and the associated “Soil”, we performed single-cell RNA sequencing (scRNA-seq) on day-18 samples from both hCOs and “Soil” (Fig. 2a). Uniform manifold approximation and projection (UMAP) analysis resolved distinct cell populations corresponding to CMs, ECs, foregut-derived cells, and CFs (Fig. 2b). Each population exhibited a characteristic set of differentially expressed genes (DEGs) (Supplementary Fig. 1a), further supported by heatmaps illustrating cluster-specific transcriptional signatures in the “Soil” (Supplementary Fig. 1b). Comparative integration of scRNA-seq datasets from hCOs and the “Soil” revealed a higher proportion of CMs in hCOs, indicated by the expression of *MYL7, MYH6*, and *MYL4*; whereas CFs, indicated by the expression of *POSTN, LMO2*, and *COL1A1*, were enriched in the “Soil” (Fig. 2c □ f; Supplementary Fig. 1c).

**Figure 2.**
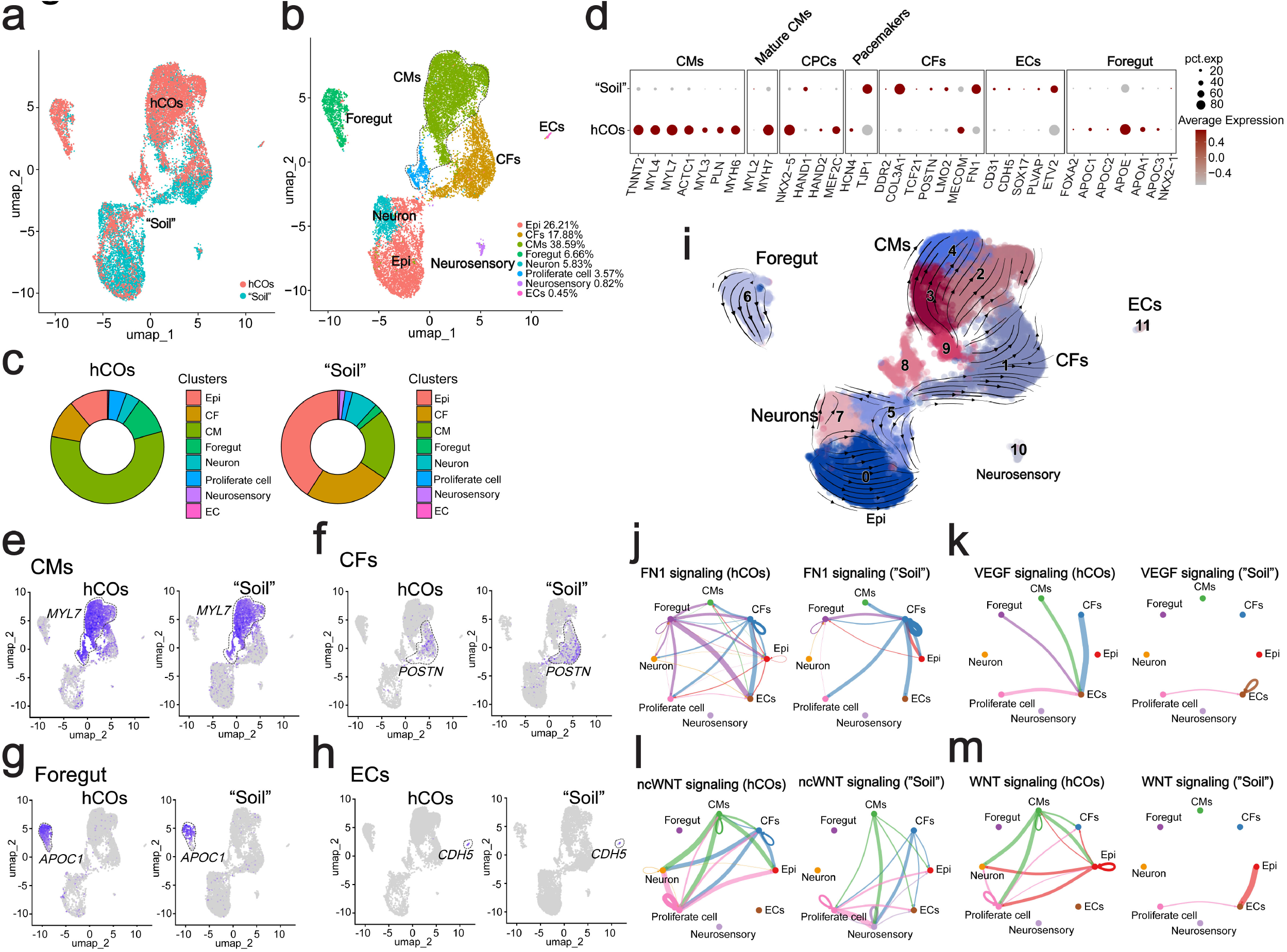
Single-cell transcriptomic profiling of hCOs and “Soil”. **a**.Integrative single-cell RNA sequencing (scRNA-seq) analysis of hCO and “Soil” samples. **b**.UMAP visualization identifying major cell clusters within hCOs and Soil. CMs, cardiomyocytes; ECs, endothelial cells; CFs, cardiac fibroblasts; Epi, epithelial cells. **c**.Pie charts showing the proportion of cell types across the two conditions. **d**.Dot plot summarizing expression of lineage markers across CM, mature CM, cardiac progenitor cell (CPC), pacemaker, CF, EC, and foregut clusters. **e**.– **f**. UMAP analyses showing enrichment of foregut and CM populations, and reduced CF populations, in hCOs compared to the “Soil” sample. **h**.Presence of ECs identified in both hCOs and “Soil” samples. **i**.RNA velocity analysis overlaid on UMAP indicating dynamic lineage trajectories. **j**.– **m**. Circle plot ligand–receptor interaction analyses highlighting differential signaling pathways in hCOs and “Soil”.

Indeed, a strong expression of foregut endoderm markers, including *APOC1, APOC2, FOXA2*, and *APOC3* were observed in both the hCOs and “Soil” (Fig. 2c, d, g; Supplementary Fig. 1c), consistent with previously reported heart/foregut models (Fig. 1c). Notably, hCOs contained a greater fraction of foregut cells than the “Soil”, suggesting their origination from heart/foregut and increasing cellular complexity within hCOs (Fig. 2c, d, f; Supplementary Fig. 1c). Together, our findings indicate that hCO formation mirrors the coordinated development of early heart/foregut structures, supporting the physiological relevance of this system for modelling early human cardiogenesis ^7^.

To define the molecular signalling networks that drive hCO self-organization, we analyzed lineage trajectories using trajectory inference and found that hCO-derived CMs arise from a distinct precursor population, cluster 9, which also goes to CFs, indicating their identity as cardiac mesodermal cells (Fig. 2i). Notably, hCOs displayed strong FN1-mediated signalling from foregut-derived cells to ECs, a feature absent in the “Soil”, suggesting that foregut signals support EC emergence (Fig. 2j). In contrast, FN1 autocrine signalling was more prominent in “Soil” CFs, consistent with fibroblast activation (Fig. 2j). In addition, hCOs exhibited robust VEGF signalling among CMs, ECs, CFs, and foregut-derived cells (Fig. 2k). Canonical and non-canonical WNT (ncWNT) signalling pathways also differed markedly between hCOs and the “Soil”, and stronger WNT-mediated signalling from CMs to neurons, proliferating cells, and epithelial cells (Epi) in hCOs (Fig. 2l, m), indicating distinct regulatory states during hCO self-organization. Together, these results define a complex intercellular signalling landscape that underlies spontaneous cardiac self-assembly from the 2D cultured hPSCs.

### Vascularized and mature hCOs are derived from the heart/foregut-like intermediate

To improve CM maturation and vascularization in hCOs, we further optimized the induction system by supplementing the differentiation medium with vascular endothelial growth factor (VEGF) at day 18 (Fig. 3a). As expected, by day 28 we observed vascularized hCOs containing abundant ECs expressing CDH5 (Fig. 3a). Analysis of the “Soil” at day 28 also revealed the presence of both CMs and ECs, with ECs closely aligned to CMs, indicating coordinated structural and functional interactions between these populations (Fig. 3b).

**Figure 3.**
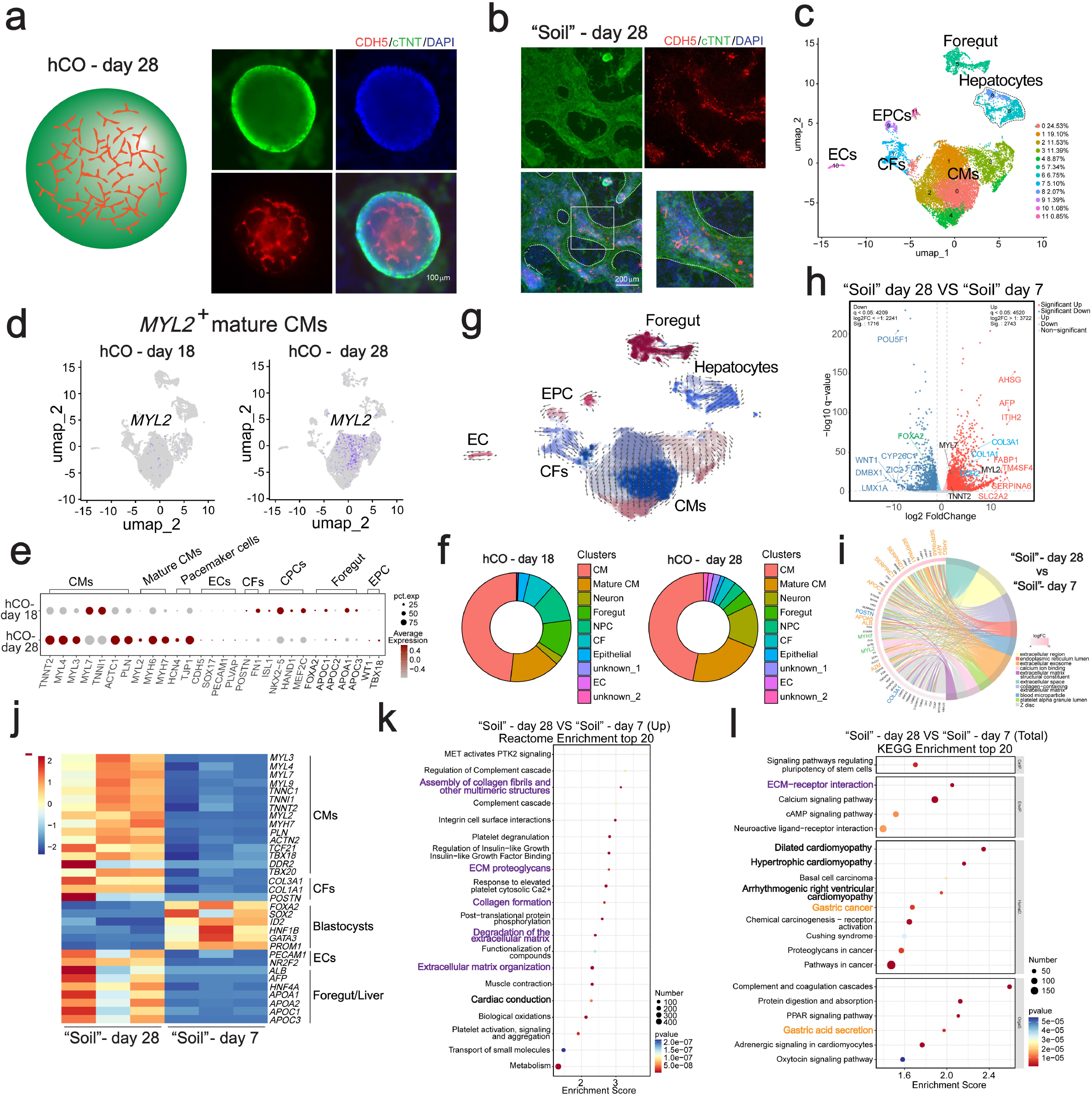
Optimized induction system promotes maturation and vascularization of hCOs. **a**.Day-28 hCOs showing extensive vascularization with abundant CDH5-positive ECs. **b**.IF analysis of “Soil” revealing close alignment of ECs and CMs. **c**.UMAP of integrated day-18 and day-28 hCO datasets identifying clusters of CMs, EPCs, ECs, CFs, and foregut-derived cells. **d**.UMAP showing increased MYL2-positive mature CMs in day-28 compared with day-18 hCOs. **e**.Dot plot summarizing lineage marker expression across CM, mature CM, CPC, pacemaker, CF, EC, and foregut clusters. **f**.Pie charts showing relative distribution of major cell types in day-18 and day-28 hCOs. **g**.RNA velocity analysis showing lineage dynamics across day-18 and day-28 hCOs. **h**.Volcano plot displaying DEGs between ““Soil”-day and “Soil”-day. **i**.Pathway enrichment analysis revealing increased regulatory activity in CM, foregut, and CF compartments between “Soil”-day 28 and “Soil”-day 7. **j**.Heatmap indicating a more mature cardiac composition in “Soil”-day 28, including enriched CMs, CFs, ECs, and foregut lineage cells. **k**– **l**. Reactome and KEGG enrichment analyses showed enrichment of cardiac, foregut, and extracellular matrix signaling pathways.

To define the cellular composition of more mature hCOs, we performed integrative scRNA-seq of day-18 and day-28 hCOs. We identified distinct clusters corresponding to CMs, EPCs, ECs, CFs, foregut cells, and hepatocytes, indicating the original heart/foregut intermediate (Fig. 3c; Supplementary Fig. 2a, b). UMAP analysis showed a significantly increased proportion of *MYL2*+CMs in day-28 hCOs (Fig. 3d), consistent with enhanced CM maturation. Dot plot and quantification analysis demonstrated increased representation of mature CMs, EPCs, CFs, and ECs in day-28 hCOs, while day-18 hCOs were enriched for foregut-derived cells, reflecting their earlier developmental stage (Fig. 3e, f; Supplementary Fig. 2a). RNA velocity analysis further indicated that mature CMs predominantly arose from immature CM populations; while EPCs transitioned toward CF lineages (Fig. 3g). These lineage dynamics mirror key aspects of *in vivo* cardiac and foregut development, supporting the physiological relevance of this system.

To evaluate the supportive role of the microenvironment, we compared bulk RNA-seq profiles of “Soil” samples at days 7 and 28. We observed higher expression of CM markers (*MYL7, TNNT2, MYL2*), hepatocyte markers (*AHSG, AFP, ITIH2, FABP1, TM4SF4, SERPINA6, SLC2A2*), CF markers (*COL3A1, COL1A1, DDR2*) in day-28 “Soil”, while foregut marker (*FOXA2*) was higher expressed in day-7 “Soil” (Fig. 3h, i). Heatmap also showed the higher expression of CM, CF, and hepatocyte markers in day-28 “Soil”, indicating their maturation (Fig. 3j). Reactome Enrichment and KEGG enrichment analysis showed significant enrichment of ECM matrix pathways, cardiac pathways, and foregut pathways in day-28 “Soil” (Fig. 3k, l). Together, these results demonstrate that the optimized hCO self-organization system recapitulates key aspects of human cardiac morphogenesis and provides a robust platform for studying heart/foregut development.

## Discussion

During early embryonic development, cardiac progenitor regions interact intimately with foregut endoderm, collaboratively driving heart tube assembly ^1^. However, the molecular and cellular mechanisms underlying this native developmental program— the “hidden magic” of organogenesis—remain poorly defined, limiting our ability to understand the cardiac self-organization and further predict congenital heart defects. Few *in vitro* models faithfully recapitulate early cardiac morphogenesis through cardiac-foregut interactions. Although current cardiac organoid systems capture key structural, functional, and cellular features, conventional iPSC-derived protocols— typically using 96-well U-bottom low-adhesion plates—fail to reproduce critical developmental events such as foregut specification and heart tube formation ^10,11^. Moreover, these approaches lack defined spatial patterning and controlled morphogen gradients in embryoid bodies, resulting in batch variability and poor reproducibility.

Current methods for producing chambered and vascularized hCOs generally depend on matrix gels and V- or U-shaped plates ^6,7,10–12^. For instance, Matrigel embedding supplies basement membrane–like ECM cues that facilitate integrin-mediated adhesion and mechano-transduction, whereas U-shaped wells impose physical confinement that enhances nutrient gradients and promotes shear-free tissue folding. Here, we describe a system for generating vascularized hCOs through transient heart/foregut-like intermediates. These heart/foregut intermediates can be applied to model Alpers–Huttenlocher syndrome harboring a POLG mutation (c.1399G>A) (Fig. 1d, e).

Despite these advances, several limitations remain. First, the generated hCOs are relatively small (50–300 µm in diameter) and typically contain a single chamber, which limits their direct translational application in cardiac regeneration. Second, hCOs at day 18 remain developmentally immature, with only a small fraction of *MYL2*+mature CMs, indicating incomplete physiological maturation. Third, due to intrinsic genetic, epigenetic, and “quality” differences among different hPSC lines, we successfully generated hCOs from human ESCs and some iPSC lines, but some iPSC lines failed to generate hCOs, which limits the widest use of this system.

## Methods

### Cardiomyocyte (CM) differentiation from human pluripotent stem cells (PSCs)

The CM differentiation protocol was adapted from previously published methods ^13^. Briefly, hPSCs were passaged at a 1:10 or 1:12 ratio and cultured for four days until reaching approximately 85% confluence. The culture medium was then replaced with **CM2**, consisting of RPMI supplemented with 1× B27 without insulin. From day 0 to day 2, **CM2** was supplemented with 6 μM CHIR99021. On day 2, the medium was refreshed with **CM2**, and on day 3, it was replaced with **CM2** supplemented with 5 μM IWR1. After 48 hours, the medium was changed back to **CM2** and subsequently refreshed every other day.

### Immunofluorescent staining

Cells were fixed in 4% paraformaldehyde (PFA) for 20 minutes at room temperature and washed three times with PBS for 5 minutes each. Samples were then incubated with permeabilization buffer, followed by three additional PBS washes. To reduce non-specific binding, samples were blocked with blocking buffer for 20 minutes at room temperature. Cells were incubated overnight at 4°C with primary antibodies diluted in staining buffer. The following day, samples were incubated with appropriate secondary antibodies for 1 h at room temperature in the dark. Fluorescence images were acquired using a fluorescence microscope.

### Flow cytometry

“Soil” samples were collected into tubes and dissociated into single-cell suspensions using Accutase (Sigma, A6964) for 10 minutes at 37°C. Cells were then fixed with 4% PFA for 10–15 minutes, washed with 500 µl FACS buffer, and centrifuged at 1,500 rpm for 5 minutes at 4°C. After carefully aspirating the supernatant, cells were permeabilized with 200 µl 0.3% Triton X-100 for 15 minutes at room temperature. Cells were washed again with 500 µl FACS buffer and centrifuged at 2,000 rpm for 5 minutes at 4°C, and the supernatant was gently removed. Cells were incubated on ice with primary antibodies for 45–60 minutes, washed with 500 µl FACS buffer, and centrifuged under the same conditions. Cells were then resuspended in 100 µl secondary antibodies, mixed thoroughly, and incubated for 20–30 minutes at room temperature in the dark. After a final wash and centrifugation, cells were resuspended in FACS buffer, filtered through 5-ml round-bottom polystyrene tubes with cell-strainer caps, and subjected to flow cytometry analysis.

### Western blot

Protein lysates were prepared by lysing hCO and “Soil” cells in RIPA buffer supplemented with protease inhibitors for 20 minutes on ice. Lysates were centrifuged at 12,000 rpm for 5 minutes at 4°C, and the supernatants were collected. Protein samples were mixed with 5× SDS sample buffer and denatured at 98°C for 10 minutes. Samples were resolved on SDS–PAGE gels, initially at 80 V and then at 120 V until the dye front reached the bottom of the gel. Polyvinylidene difluoride (PVDF) membranes were equilibrated in transfer buffer for 1–2 minutes, briefly rinsed, and assembled with the gel in a transfer apparatus. Proteins were transferred at 110 V for 75 minutes. Membranes were then blocked in blocking buffer for 1 h at room temperature. Primary antibodies diluted in blocking buffer were incubated with the membranes overnight at 4°C. After three washes with TBST for 10 minutes each, membranes were incubated with HRP-conjugated secondary antibodies diluted in blocking buffer for 1 h at room temperature, followed by three additional 10-minute washes with TBST. The membranes were then ready for imaging and further analysis.

### Real-time PCR Analysis

Total RNA was extracted from collected cells using TRIzol reagent (Invitrogen) according to the manufacturer’s instructions. Approximately 400 ng of RNA was reverse-transcribed into complementary DNA (cDNA). Quantitative PCR (qPCR) was performed using SYBR Green chemistry, and relative mRNA expression levels were calculated using the ΔΔCt method. *GAPDH* was selected as the endogenous reference gene for normalization. All qPCR assays were conducted with technical replicates to ensure reproducibility.

### Image processing and data quantification

IF images were analyzed using custom macros in ImageJ. Three-dimensional (3D) reconstruction and volumetric analysis of hCOs were performed using the 3D ImageJ Suite.

### ScRNA-seq for hCOs and “Soil”

hCOs at day 12, day 18, day 28 and Soil sample at day 18 were collected and dissociated into single cells using Accutase. The dissociated cells were washed with PBS and centrifuged for 5 minutes; the cells were resuspended ensuring that there were above 100 × 10^3^ cells for each sample. Library preparation for scRNA-seq analysis of the mRNA was performed according to the Chromium NextGem Single Cell 3□ Reagent Kits v3.1 user guide. Briefly, a given excess of cells was loaded to the 10x controller to reach a target number of around 1□×□10^4^□a given excess. Quantification of the libraries was performed using Qubit 4 Fluorometer.

ScRNA-seq was conducted using the NextSeq 2000 platform, following the manufacturer’s protocol. In brief, a single-cell suspension was loaded into the microfluidic chip together with gel beads, reverse transcription reagents, and oil. The microfluidic system partitioned the mixture into thousands to tens of thousands of Gel Beads-in-Emulsion (GEMs), each ideally encapsulating a single cell and a uniquely barcoded bead. Within each GEM, cells were lysed, and mRNA transcripts were captured by the bead-bound barcoded primers. Reverse transcription was subsequently performed inside the GEMs to synthesize complementary cDNA from the captured mRNA. Following reverse transcription, the GEMs were broken to release the barcoded cDNA, which was then amplified by PCR to generate sufficient material for downstream processing. The amplified cDNA was used to construct sequencing libraries compatible with Illumina sequencing platforms through the addition of sequencing adapters. Finally, the prepared libraries were sequenced on Illumina instruments, producing paired-end reads containing both cell barcodes and transcript sequences. The resulting RNA sequence data were aligned to the human reference genome for further analysis.

### ScRNA-seq raw data processing

Single-cell RNA-seq data were processed using the 10x Genomics Cell Ranger pipeline. Binary base call (BCL) files were demultiplexed into FASTQ files using the cellranger mkfastq command with the appropriate sample sheet and barcode sets. Reads were processed with Cell Ranger (version 8.0.1) using default parameters for demultiplexing, alignment to the GRCh38 reference genome, and generation of gene–cell count matrices. Quality control was performed in R using Seurat (version 5.3.1). Cells with fewer than 500 detected genes or with mitochondrial gene content >10% were excluded.

Gene expression matrices were normalized using the LogNormalize method. Principal component analysis (PCA) was performed, followed by dimensionality reduction with UMAP. Clustering was conducted using the Louvain algorithm. Two-dimensional UMAP plots were generated using the DimPlot function in Seurat. Cluster marker genes were identified using the FindMarkers function with default parameters. For each cluster, the top 10 marker genes were used to annotate cell types, and a heatmap of these genes was generated using the DoHeatmap function. Dot plots for selected genes of interest across clusters were generated using the DotPlot function, where dot size represents the percentage of expressing cells and dot color reflects mean expression level.

### CellChat Analysis of Cell–Cell Communication

Intercellular communication networks were inferred using the CellChat R package (version 2.2.0). Normalized gene expression matrices and cell-type annotations served as input. CellChat uses a curated database of ligand–receptor interactions to predict putative signaling events between cell populations. Significant signaling pathways were identified using CellChat’s built-in statistical framework. Pathway enrichment and comparisons between hCO and “Soil” samples were performed to highlight differences in intercellular communication. Cell–cell interaction networks for significant pathways were visualized using the netVisual_aggregate function, where each node represents a cell cluster, line segments represent interactions, and arrows indicate the direction of signaling.

### RNA velocity analysis

Differentiation trajectories of hCO, “Soil,” and mature hCO cell clusters were inferred using RNA velocity. Spliced and unspliced reads were quantified with velocyto (v0.17.17) using default settings. Downstream analyses and visualizations were performed separately for each dataset using scVelo (v0.2.4) in Python. Marker gene expression across clusters was visualized with violin plots generated using the VlnPlot function in Seurat, illustrating the distribution of marker expression within each cell population.

### Bulk RNA-seq library prep

For bulk RNA-seq, total RNA was isolated from hPSCs, early hCOs, late hCOs, and CMs using TRIzol (Invitrogen) according to the manufacturer’s protocol. Libraries were prepared by Ouyi Biomedical Technology Co., Ltd. (China) using the NEBNext Ultra II RNA Library Prep Kit (New England Biolabs, E7805S). The resulting libraries had fragment sizes of 220–500 bp, with a peak around 350 bp. Sequencing was performed on a BGISEQ-500 platform to generate 150-bp paired-end reads. Differentially expressed genes (DEGs) were initially identified using thresholds of expression percentage >0.1, log2 fold change >0.25, and adjusted p-value below the chosen significance cutoff. Volcano plots were generated in R using the geom_point function in ggplot2. Principal component analysis (PCA) was performed using the getDimRedPlot function from the muRtools R package (v0.9.5), based on the first two principal components.

### Bulk RNA-seq data analysis

Raw sequencing reads were subjected to quality control with FastQC. Clean reads were aligned to the GRCh38 reference genome using bowtie2 (version 2.2.5). Alignment quality was assessed with SAMtools. Gene-level read counts were obtained using in-house scripts. Normalization and differential expression analysis were performed using DESeq2 in R. Genes with adjusted p-value <0.05 and |log2 fold change| ≥1 were considered significantly differentially expressed. Significantly regulated genes were subjected to Gene Ontology (GO) and Kyoto Encyclopedia of Genes and Genomes (KEGG) enrichment analysis using the clusterProfiler R package, and enrichment results were visualized using ggplot2.

### Statistics and reproducibility

At least three independent experimental replicates were performed for each condition, with each replicate conducted in triplicate. Statistical analyses were carried out using GraphPad Prism software (version 6). Data are presented as mean ± standard deviation (SD). The sample size (n) corresponds to the number of independent biological replicates. Statistical significance between groups was assessed using an unpaired Student’s *t*-test. A threshold of *p* < 0.05 was considered statistically significant. Significance levels are denoted as follows: ns, not significant (*p* > 0.05); \**p* < 0.05; \*\**p* < 0.01; and \*\*\**p* < 0.001.

## Acknowledgements

This work was supported by a Natural Science Foundation of China (NSFC)-Young Scientists Fund (#82200314), Innovation and Technology Commission of the Hong Kong Government grants (GHX09721SZ and MHP00523), Direct Grant for Research (#2021.069 and #2022.082) from the Chinese University of Hong Kong (CUHK), the National Natural Science Foundation of China (NSFC)/Research Grants Council (RGC) Joint Research Scheme (#N_CUHK428/22), and Strategic Seed Funding for Collaborative Research Scheme (#MK/WW/SSFCRS2324/8014/24jh) from CUHK, RGC general research fund (#14116825), the State Ministries Special Budget to support the MOE Key Laboratory for Regenerative Medicine (CUHK–Jinan University) [2622009], partially supported by Science and Technology Planning Project of Guangdong Province (2023B1212120009), Guangdong Basic and Applied Basic Research Fund (2023B0303000023), Co-funding Mechanism on Joint Laboratories with the CAS (JLFS/M-403/24), CUHK–Shandong University Joint Laboratory on Reproductive Genetics, and CUHK Laboratory Support Special Fund for Key Laboratory for Regenerative Medicine, Ministry of Education, China. This work was supported (in full or partially) by the research fund to the Center for Neuromusculoskeletal Restorative Medicine from Health@InnoHK program launched by Innovation and Technology Commission, Hong Kong.

## Author Contributions

W.W. performed the majority of the experiments. W.W., G.L., and W.-Y.C. designed the experiments. Q.H. and H.Z. carried out cell culture and molecular assays. Z.X. and L.Y.Y. provided bioinformatics support. X.S. and Q.M. performed scRNA-seq, and Y.G., Y.F., X.L., and X.B. contributed to data analysis. G.L. and W.-Y.C. conceived the study. W.W., G.L., and W.-Y.C. interpreted the data and wrote the manuscript, with input from all authors.

## References

1. Hosseini, H.S., Garcia, K.E., and Taber, L.A. (2017). A new hypothesis for foregut and heart tube formation based on differential growth and actomyosin contraction. Development 144, 2381–2391. 10.1242/dev.145193.

2. Aleksandrova, A., Czirok, A., Kosa, E., Galkin, O., Cheuvront, T.J., and Rongish, B.J. (2015). The endoderm and myocardium join forces to drive early heart tube assembly. Dev Biol 404, 40–54. 10.1016/j.ydbio.2015.04.016.

3. Peng, T., Tian, Y., Boogerd, C.J., Lu, M.M., Kadzik, R.S., Stewart, K.M., Evans, S.M., and Morrisey, E.E. (2013). Coordination of heart and lung co-development by a multipotent cardiopulmonary progenitor. Nature 500, 589– 592. 10.1038/nature12358.

4. Tiburcy, M., Hudson, J.E., Balfanz, P., Schlick, S., Meyer, T., Chang Liao, M.L., Levent, E., Raad, F., Zeidler, S., Wingender, E., et al. (2017). Defined Engineered Human Myocardium With Advanced Maturation for Applications in Heart Failure Modeling and Repair. Circulation 135, 1832–1847. 10.1161/CIRCULATIONAHA.116.024145.

5. Mills, R.J., Titmarsh, D.M., Koenig, X., Parker, B.L., Ryall, J.G., Quaife-Ryan, G.A., Voges, H.K., Hodson, M.P., Ferguson, C., Drowley, L., et al. (2017). Functional screening in human cardiac organoids reveals a metabolic mechanism for cardiomyocyte cell cycle arrest. Proc Natl Acad Sci U S A 114, E8372–E8381. 10.1073/pnas.1707316114.

6. Schmidt, C., Deyett, A., Ilmer, T., Haendeler, S., Torres Caballero, A., Novatchkova, M., Netzer, M.A., Ceci Ginistrelli, L., Mancheno Juncosa, E., Bhattacharya, T., et al. (2023). Multi-chamber cardioids unravel human heart development and cardiac defects. Cell 186, 5587–5605 e5527. 10.1016/j.cell.2023.10.030.

7. Drakhlis, L., Biswanath, S., Farr, C.-M., Lupanow, V., Teske, J., Ritzenhoff, K., Franke, A., Manstein, F., Bolesani, E., and Kempf, H. (2021). Human heartforming organoids recapitulate early heart and foregut development. Nature Biotechnology 39, 737–746.

8. Silva, A.C., Matthys, O.B., Joy, D.A., Kauss, M.A., Natarajan, V., Lai, M.H., Turaga, D., Blair, A.P., Alexanian, M., Bruneau, B.G., and McDevitt, T.C. (2021). Co-emergence of cardiac and gut tissues promotes cardiomyocyte maturation within human iPSC-derived organoids. Cell Stem Cell 28, 2137–+. 10.1016/j.stem.2021.11.007.

9. Rahman, S., and Copeland, W.C. (2019). POLG-related disorders and their neurological manifestations. Nat Rev Neurol 15, 40–52. 10.1038/s41582-018-0101-0.

10. Hofbauer, P., Jahnel, S.M., Papai, N., Giesshammer, M., Deyett, A., Schmidt, C., Penc, M., Tavernini, K., Grdseloff, N., and Meledeth, C. (2021). Cardioids reveal self-organizing principles of human cardiogenesis. Cell 184, 3299–3317.e3222.

11. Abilez, O.J., Yang, H., Guan, Y., Shen, M., Yildirim, Z., Zhuge, Y., Venkateshappa, R., Zhao, S.R., Gomez, A.H., El-Mokahal, M., et al. (2025). Gastruloids enable modeling of the earliest stages of human cardiac and hepatic vascularization. Science 388, eadu9375. 10.1126/science.adu9375.

12. Schmidt, C., Deyett, A., Ilmer, T., Caballero, A., Haendeler, S., Pimpale, L., Netzer, M., Ginistrelli, L., Cirigliano, M., and Mancheno, E. (2022). Multichamber cardioids unravel human heart development and cardiac defects. bioRxiv 2022.07.14.499699.

13. Burridge, P.W., Matsa, E., Shukla, P., Lin, Z.C., Churko, J.M., Ebert, A.D., Lan, F., Diecke, S., Huber, B., Mordwinkin, N.M., et al. (2014). Chemically defined generation of human cardiomyocytes. Nat Methods 11, 855–860. 10.1038/nmeth.2999.

