## Supplementary Figures for "Vascularized Organoids Emerge from a Heart–Foregut-like Intermediate Recapitulating Early Human Development"

**Figure S1. ScRNA-seq analysis of the "Soil-Seed" system.**
**a**. Dot plot summarizing lineage-specific marker gene expression for CMs, mature CMs, CPCs, CFs, and foregut-derived cells.
**b**. Heatmap displaying gene expression profiles across distinct cell lineages.
**c**. UMAP visualization showing clustering of different cell populations based on lineage-specific marker genes.

**Figure S2. ScRNA-seq analysis of mature hCOs.**
**a**. UMAP plots comparing cell populations in day 18 and day 28 hCOs.
**b**. Characterization of distinct clusters based on signature marker genes.
**c**. KEGG pathway enrichment analysis highlighting ECM–receptor interaction and cardiac signaling pathways.

### Figure S1.

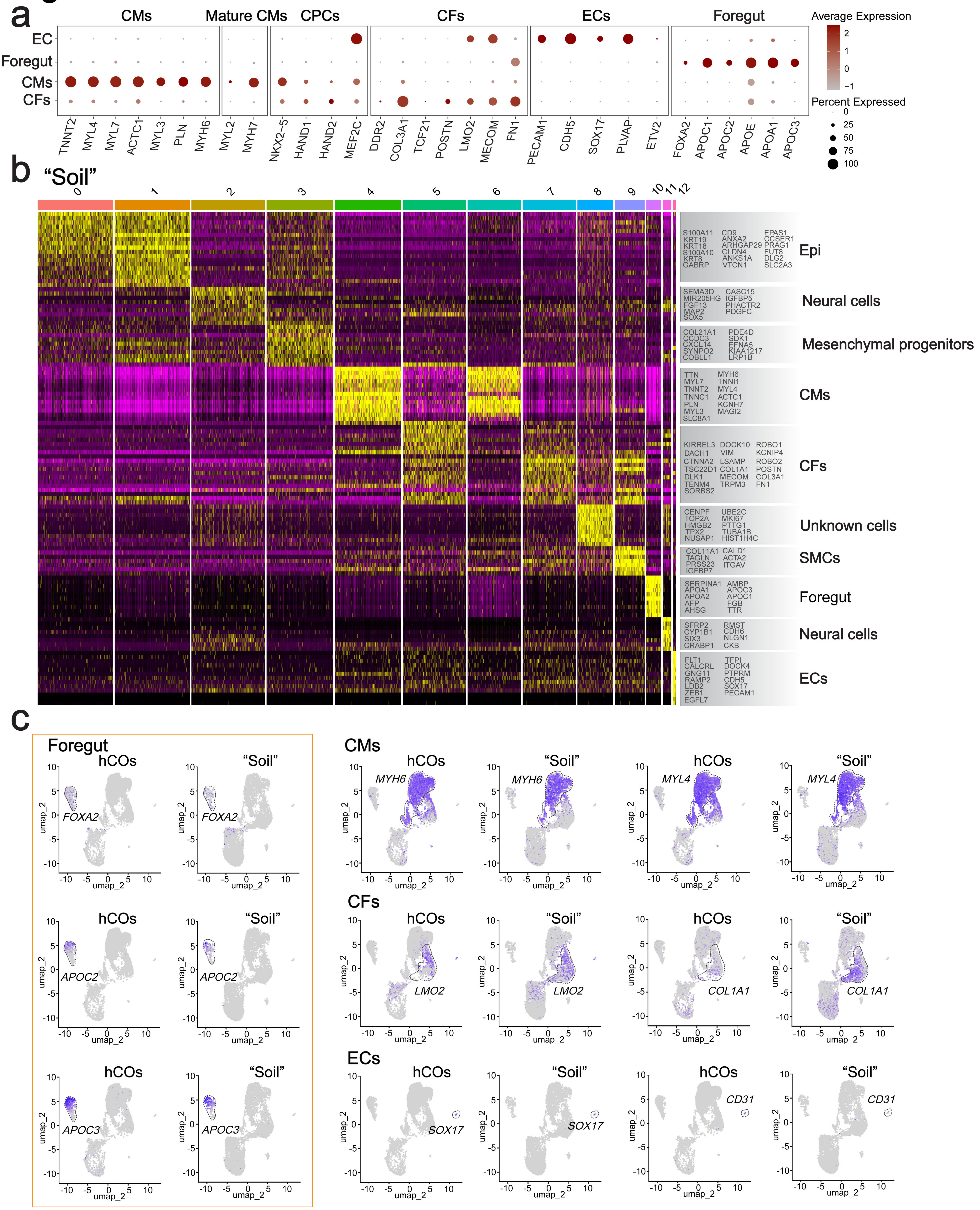
a
CMs
Mature CMs
CPCs
CFs
ECs
Foregut
EC
Foregut
CMs
CFs
TNNT2
MYL4
MYL7
ACTC1
MYL3
PLN
MYH6
MYL2
MYH7
NKX2-5
HAND1
HAND2
MEF2C
DDR2
COL3A1
TCF21
POSTN
LMO2
MECOM
FN1
PECAM1
CDH5
SOX17
PLVAP
ETV2
FOXA2
APOC1
APOC2
APOE
APOA1
APOC3
Average Expression
Percent Expressed
b
"Soil"
0
1
2
3
4
5
6
7
8
9
10
11
12
S100A11 KRT19 KRT18 S100A10 KRT8 GABRP CD9 ANXA2 ARHGAP29 CLDN4 ANKS1A VTCN1 EPAS1 CCSER1 PRAG1 FUT8 DLG2 SLC2A3
Epi
SEMA3D MIR205HG FGF13 MAP2 SOX5 CASC15 IGFBP5 PHACTR2 PDGFC
Neural cells
COL21A1 CCDC3 CXCL14 SYNPO2 COBLL1 PDE4D SDK1 EFNA5 KIAA1217 LRP1B
Mesenchymal progenitors
TTN MYL7 TNNT2 TNNC1 PLN MYL3 SLC8A1 MYH6 TNNI1 MYL4 ACTC1 KCNH7 MAGI2
CMs
KIRREL3 DACH1 CTNNA2 TSC22D1 DLK1 TENM4 SORBS2 DOCK10 VIM LSAMP COL1A1 MECOM TRPM3 ROBO1 KCNIP4 ROBO2 POSTN COL3A1 FN1
CFs
CENPF TOP2A HMGB2 TPX2 NUSAP1 UBE2C MKI67 PTTG1 TUBA1B HIST1H4C
Unknown cells
COL11A1 TAGLN PRSS23 IGFBP7 CALD1 ACTA2 ITGAV
SMCs
SERPINA1 APOA1 APOA2 AFP AHSG AMBP APOC3 APOC1 FGB TTR
Foregut
SFRP2 CYP1B1 SIX3 CRABP1 RMST CDH6 NLGN1 CKB
Neural cells
FLT1 CALCRL GNG11 RAMP2 LDB2 ZEB1 EGFL7 TFPI DOCK4 PTPRM CDH5 SOX17 PECAM1
ECs
c
Foregut
hCOs
"Soil"
FOXA2
APOC2
APOC3
CMs
MYH6
MYL4
CFs
LMO2
COL1A1
ECs
SOX17
CD31
umap_2

### Figure S2.

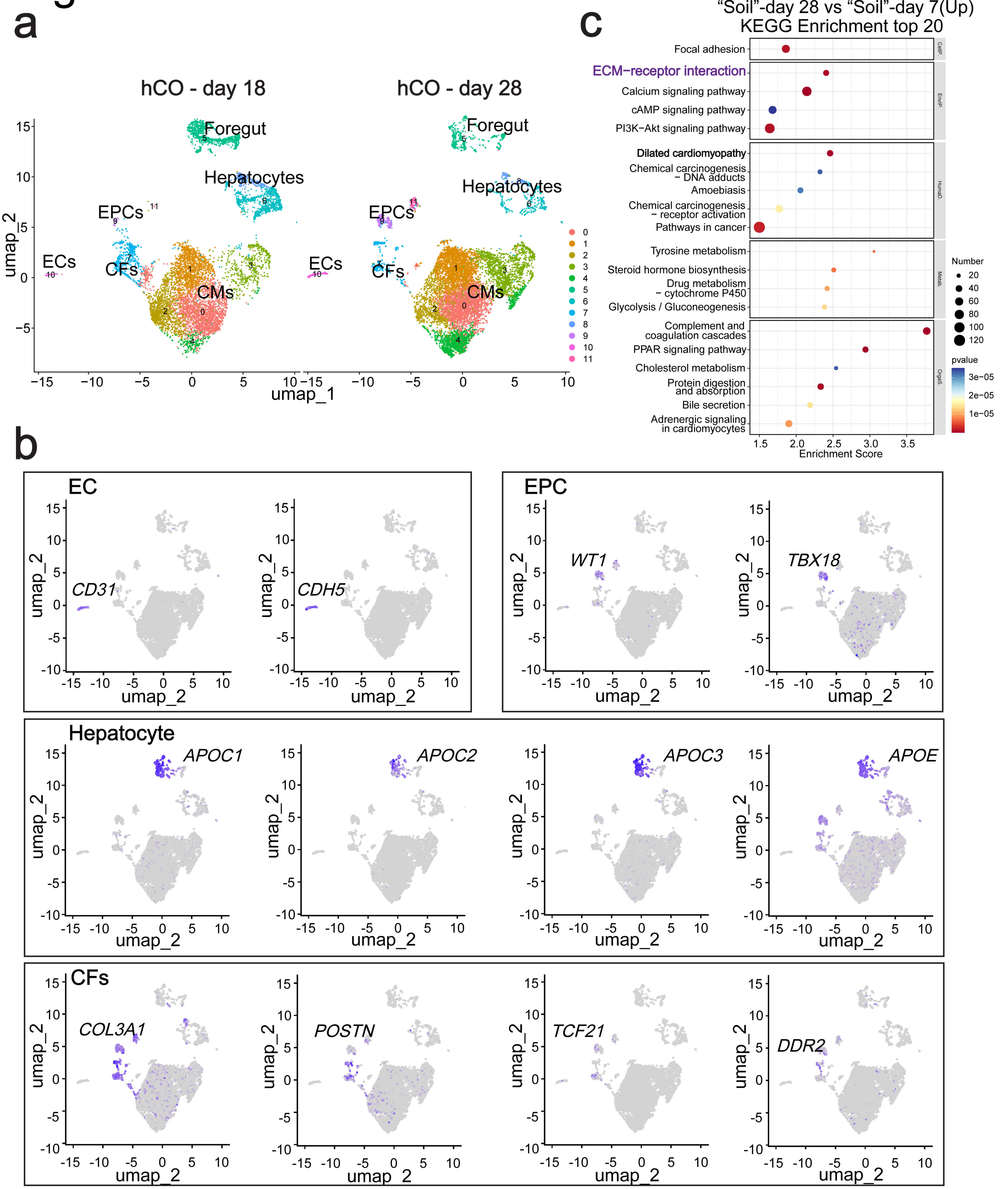
a
hCO - day 18
hCO - day 28
Foregut
Hepatocytes
EPCs
ECs
CFs
CMs
umap_1
umap_2
c
“Soil”-day 28 vs “Soil”-day 7(Up)
KEGG Enrichment top 20
Focal adhesion
ECM−receptor interaction
Calcium signaling pathway
cAMP signaling pathway
PI3K−Akt signaling pathway
Dilated cardiomyopathy
Chemical carcinogenesis − DNA adducts
Amoebiasis
Chemical carcinogenesis − receptor activation
Pathways in cancer
Tyrosine metabolism
Steroid hormone biosynthesis
Drug metabolism − cytochrome P450
Glycolysis / Gluconeogenesis
Complement and coagulation cascades
PPAR signaling pathway
Cholesterol metabolism
Protein digestion and absorption
Bile secretion
Adrenergic signaling in cardiomyocytes
Enrichment Score
Number
pvalue
b
EC
CD31
CDH5
EPC
WT1
TBX18
Hepatocyte
APOC1
APOC2
APOC3
APOE
CFs
COL3A1
POSTN
TCF21
DDR2
